# SnapperDB: A database solution for routine sequencing analysis of bacterial isolates

**DOI:** 10.1101/189118

**Authors:** Timothy Dallman, Philip Ashton, Ulf Schafer, Aleksey Jironkin, Anais Painset, Sharif Shaaban, Hassan Hartman, Richard Myers, Anthony Underwood, Claire Jenkins, Kathie Grant

## Abstract

Real-time surveillance of infectious disease using whole genome sequencing data poses challenges in both result generation and communication. SnapperDB represents a set of tools to store bacterial variant data and facilitate reproducible and scalable analysis of bacterial populations. We also introduce the ‘SNP address’ nomenclature to describe the relationship between isolates in a population to the single nucleotide resolution.

**Summary:** We announce the release of SnapperDB v1.0 a program for scalable routine SNP analysis and storage of microbial populations.

**Availability:** SnapperDB is implemented as a python application under the open source BSD license. All code and user guides are available at https://github.com/phe-bioinformatics/snapperdb.

**Supplementary information:** Supplementary data are available at Bioinformatics online.

## 1 Introduction

The As routine whole genome sequencing (WGS) of bacterial isolates for infectious disease surveillance becomes a reality ^1–3^ scalable data storage solutions are required. Analysis of bacterial populations often requires re-computing the genomic variants across all relevant isolates which is not feasible in rapidly growing, large datasets. Public Health England has embarked on the implementation of high throughput, realtime sequencing for the surveillance of several important human pathogens and aims to leverage the high discriminatory power of single nucleotide polymorphisms (SNPs) to detect linked cases and outbreaks of infectious disease. In this application note we present SnapperDB, a set of tools to store bacterial variant data to facilitate reproducible and scalable analysis of bacterial populations. We introduce the ‘SNP address’ nomenclature to describe the relationship between isolates in a population to the nucleotide resolution.

## 2 Features

The Variant calling in bacterial genomics generally relies on mapping short reads to a single reference genome and this is the central tenet of SnapperDB. The input to SnapperDB is either (1) sequence data in FASTQ format and a reference genome or (2) a Variant Call Format (VCF) file generated for each strain with all positions emitted. If sequence data is provided, SnapperDB utilises PHEnix (https://github.com/phe-bioinformatics/PHEnix) to execute third party mapping (e.g. BWA^4^, Bowtie^5^) and variant calling (e.g. GATK^6^, MPileup^7^) software of the users choice. This VCF is parsed with user specified thresholds of mapping quality, mapping depth and variant ratio to identify those positions that are of high quality. Polymorphisms that do not meet these criteria, positions that have no aligned reads, or invariant positions with depth or mapping quality less than the specified thresholds are termed “ignored positions”.

For each strain, SnapperDB stores the variant positions and the ignored positions in a PostgreSQL database, with positions the same as the reference genome not stored. If the reference genome is provided in GenBank format, for each variant detected a set of characteristics about that variant is also stored (e.g. coding/non-coding, gene location, synonymous/non-synonymous, pseudogene etc). One utility of SnapperDB is the output of high quality variant positions for a user-defined selection of strains in FASTA format for phylogenetic inference. Options to output whole genome alignments, or alignments that employ partial or complete deletion of missing positions are available. SnapperDB is compatible with the GFF output from recombination detection software such as Gubbins^8^ to mask positions from alignments as required.

**Fig. 1.**
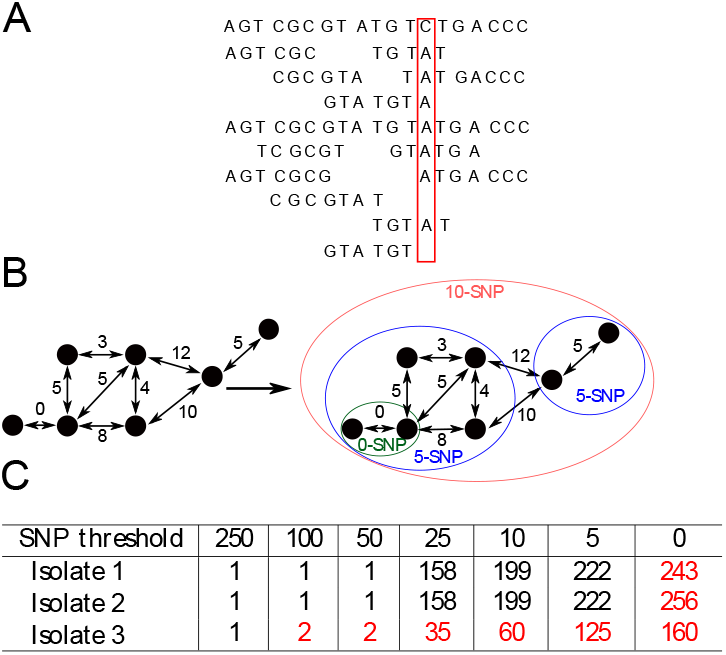
**A. Illustration of SNP difference between a reference sequence (top row) and a set of isolate sequences (remaining row). B. Single linkage clustering of SNP differences into 0, 5 and 10 SNP levels for a set of isolates. C. Examples of SNP addresses based on the seven descending SNP thresholds; 250, 100, 50, 25, 10, 5, 0**.

As new strains are added to SnapperDB they are compared to the database and a distance matrix is maintained of pairwise SNP distances. Furthermore, this matrix is available for clustering. Single linkage clustering of genetic distance is an effective method of describing phylogenetic groups as it is inclusive of clonal expansion events. Using hierarchical single linkage clustering of pairwise SNP distances we can derive an isolate level nomenclature for each genome sequence allowing efficient searching of the population studied as well as facilitating automated, real-time cluster detection. Single linkage clustering is performed at seven descending thresholds of SNP distance; 250, 100, 50, 25, 10 and 0 SNPs. This clustering results in a discrete seven-digit code where each number represents the cluster membership at each descending SNP distance threshold. The resultant ‘SNP address’ provides an isolate level nomenclature where two isolates with the same SNP addresses have 0 SNP differences. The SNP address has become the primary whole genome sequencing result for the surveillance of food-borne pathogens in the England and has been utilised as the case definition in international outbreak investigations.

## 3 Conclusions

Real-time surveillance of infectious disease using WGS data poses challenges in both result generation and communication. In this article, we introduce SnapperDB an end-to-end solution for processing of FASTQ reads to high quality variants that can be stored in a queryable database. This application has enabled Public Health England to embark on whole genome sequencing pathogen surveillance at a national level for *Salmonella, E. coli, Shigella* and *Listeria* sequencing over 20,000 genomes over since 2014. It has enabled us to track persistence sources of *Salmonellosis*^9,10^ and facilitated several investigations into outbreaks of Shiga Toxin producing *E. coli* (STEC)^11–13^.

The SNP address is the first strain level nomenclature, to our knowledge, developed for WGS pathogen surveillance and provides a framework for cluster identification, definition and investigation.

